# Functional phenotyping of small multidrug resistance proteins from *Staphylococcus aureus* and *Francisella tularensis* reveals functional homology to EmrE

**DOI:** 10.1101/2021.01.06.425668

**Authors:** Peyton J. Spreacker, Will F. Beeninga, Brooke L. Young, Colin J. Porter, Katherine A. Henzler-Wildman

## Abstract

Small multidrug resistance (SMR) transporters efflux toxic substrates from bacterial cells. These transporters were recently divided into two subfamilies: the GdX-like and EmrE-like SMRs. The EmrE-like subfamily of SMRs is predicted to contain transporters that are highly promiscuous in both substrate specificity and mechanism based on extensive characterization of the founding member of this subfamily, EmrE. However, there is only limited functional analysis of other members of this family from pathogenic strains such as *Staphylococcus aureus* and *Francisella tularensis*. Here, we use a small compound screen to explore the substrate specificity and diversity of EmrE-subfamily SMRs from these two bacterial species and confirm that they are functionally more like EmrE than the GdX-like subfamily of toxic-metabolite transporters. The results of these experiments lay the foundation for understanding the complex substrate specificity profiles of SMR family transporters and assess the potential for targeting these transporters for future antibiotic development, either broadly or in a species-specific manner.

## INTRODUCTION

Multidrug resistance (MDR) transporters actively efflux antibiotics, antiseptics, and other toxic compounds, reducing intracellular concentrations sufficiently to enable bacterial survival. The small multidrug resistance (SMR) transporters are the smallest known active transporters and are found only in bacteria. In addition to toxin efflux, several studies have implicated SMR transporters in bacterial motility, virulence, and biofilm formation, although how SMR transport activity contributes to these functions is not yet fully understood (1–5). The *E. coli* transporter, EmrE, was the first SMR transporter to be functionally characterized (6), both *in vitro* and *in vivo*, and is still the most extensively studied SMR homolog. EmrE functions as an antiparallel homodimer and is a highly promiscuous transporter that exports polyaromatic cations and quaternary ammonium cations across the inner membrane of *E. coli* (5–9). The well-established antiport of 2 H^+^/1 substrate by EmrE enables active efflux of substrate by coupling it to inward movement of protons down the proton motive force (6, 10–13). This antiport activity leads to the well-established ability of EmrE to confer resistance to toxic cations.

Members of the SMR family are readily identified through sequence analysis (14–16) and have been divided into two subclasses: EmrE-like small multidrug pumps with broad substrate specificity and GdX (formerly SugE) -like transporters have narrower substrate specificity profile and confer resistance to a smaller set of toxic cations. Recently, the GdX-like subfamily was reclassified as toxic metabolite exporters (15, 17) as a result of the discovery that expression of this subfamily is regulated by a novel family of riboswitches (16, 18, 19) in response to guanidinium^+^. Furthermore, transport assays confirmed efficient 2H^+^/1 guanidinium^+^ antiport and selectivity for guanidinium over other closely related compounds by GdX (15). However, there may be exceptions the definition of the GdX subfamily as more selective, tightly coupled metabolite exporters and the EmrE subfamily as highly promiscuous exporters of exogenous toxins. For example, MdtIJ, a heterodimer classified as EmrE-like via sequence, was shown to transport spermidine, a metabolite, rather than acting as a multidrug transporter (20, 21). This raises the question of how much functional variation exists within each subfamily, and how well functional phenotypes correlate with bioinformatic classifications, since experimental assessment of transport or drug resistance phenotypes has only been performed for a relatively small number SMR homologs and limited number of substrates (1, 9, 15, 22–26). Here we experimentally confirm the EmrE-like classification of two SMR homologs from bacterial pathogens, SMR_Sa_ from *Staphylococcus aureus* and SMR_Ft_ from *Francisella tularensis* by comparing the results of resistance assays of these SMRs to EmrE. We also determine the substrate promiscuity of EmrE-like SMRs by performing diverse compound screens against EmrE, SMR_Sa_, and SMR_Ft_.

The *E. coli* SMR transporter EmrE has become a model system for biophysical studies of proton-coupled transport. Throughout several decades of research on EmrE, new data has continued to reveal surprising features of EmrE structure and mechanism and to refine our understanding of its transport activity. EmrE has been implicated in resistance to not only quaternary ammonium cations and other toxins, but also in increasing bacterial survival in the presence of environmental and osmotic stress (1). EmrE has also been tied to bacterial processes such as biofilm formation (5). Recent mechanistic work has demonstrated that EmrE is not a strictly coupled proton/drug antiporter in the manner of the GdX-subfamily and may be able to perform uniport and symport (12) in addition to antiport. These other modes of transport have major biological implications (Figure 1). Compared to the resistance conferred by antiport of drug and proton (Figure 1A), symport would result in influx of toxic substrates alongside protons and would result in susceptibility rather than resistance (Figure 1B). Uncoupled transport of either proton or positively charged toxic substrates would similarly result in influx of proton or toxin due to the inwardly directed proton-motive force and negative-inside membrane potential in bacteria. Thus, these transport modes would also result in susceptibility due to rundown of the proton motive force or toxin uptake (27). Prior experiments showed that substrate identity determines the rate of alternating access and efflux rates in bacteria (28). More recent kinetic modeling illustrates how different substrates can potentially switch the transport behavior from antiport to symport, or uniport through drug-dependent differences in the rate constants for key steps in the transport cycle, such as substrate off-rate, alternating access rate of substrate-bound transporter or altered protonation and deprotonation rates when drug is bound (27). The mechanistic promiscuity suggested by the biophysical data (12) and kinetic simulations (27), increases the importance of understanding what defines an SMR substrate and whether transport direction (influx versus efflux) and biological outcome (resistance versus susceptibility) vary based on the identity of the substrate itself.

**Figure 1:**
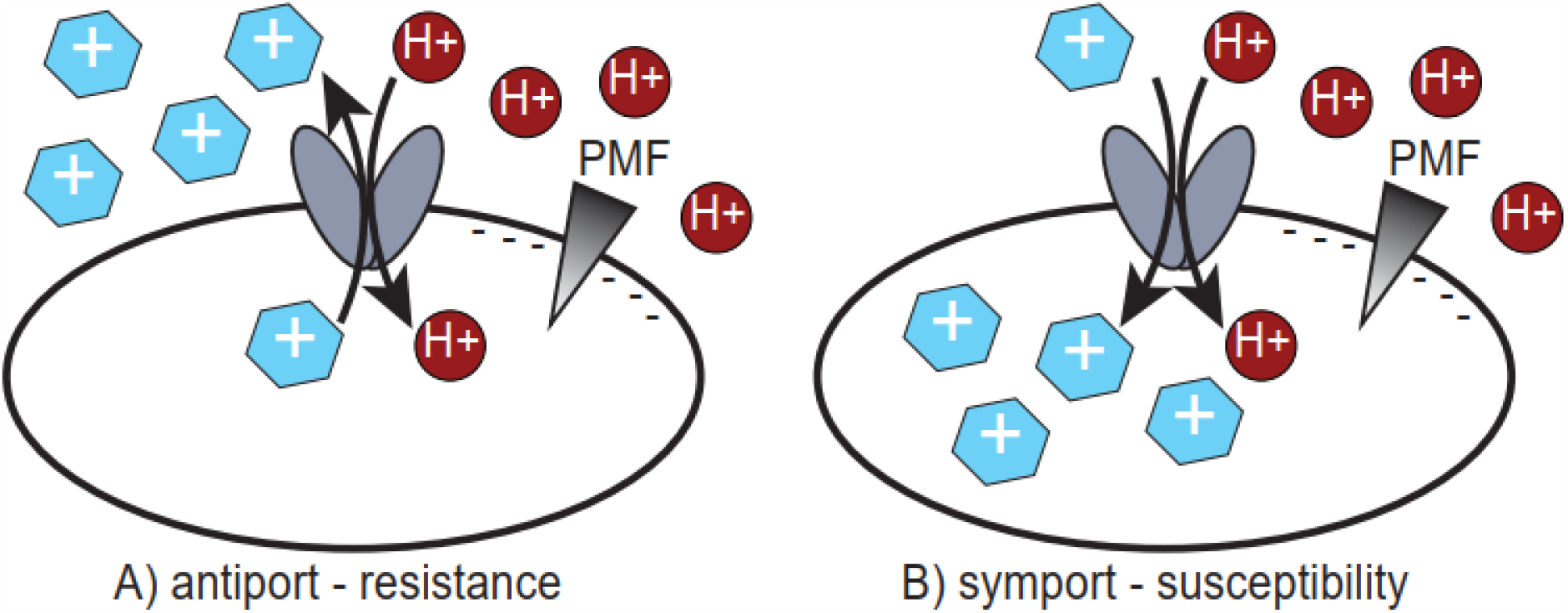
Different transport modes in EmrE-like SMRs have important biological implications. Cartoon representations of the different transport modes possible in EmrE-like SMRs with their corresponding biological phenotype. Antiport confers resistance to the substrate. Symport and drug or proton uniport confer susceptibility to the given substrate.

We functionally characterized EmrE as well as the EmrE-like SMR homologs SMR_Sa_ and SMR_Ft_ using Biolog assays to profile the metabolic activity of *E. coli* expressing either WT or non-funtional variant in the presence of 240 different compounds. EmrE has only been screened for resistance against a few dozen compounds (9, 11, 15, 22, 25, 26), so this will broaden our knowledge of what is and is not an EmrE substrate. SMR_Sa_ and SMR_Ft_ proteins were selected for comparison as they are present in human pathogens and are likely to be relevant to drug resistance in their respective pathogens. Despite their presence in diverse bacteria, all three proteins are bioinformatically similar, with 42-44% similarity between EmrE and its homologs from *S. aureus* and *F. tularensis*. Our assays characterize *both* substrate resistance *and* susceptibility, assess substrate promiscuity for each transporter, and compare the substrate profiles for these three SMR homologs. We find that SMR_Sa_ and SMR_Ft_ are functionally like EmrE: promiscuous transporters with many substrates, and that they can confer either resistance or susceptibility to toxic compounds. This experimentally demonstrates that promiscuity in both substrate and mechanism apply more broadly to the EmrE-like SMR subfamily, including SMR homologs from clinical pathogens.

## RESULTS

### Characterization of GdX-like versus EmrE-like behavior using representative substrates

EmrE-like behavior is characterized by the ability to confer resistance to broad classes of quaternary ammonium compounds and polyaromatic cations, such as methyl viologen (MV^2+^) and ethidium (Eth^+^), but not smaller metabolites like guanidinium (1, 11, 15). GdX-like activity is characterized by a narrower substrate specificity and the ability to confer resistance to guanidinium but not MV^2+^ or Eth^+^(15). We therefore used microplate growth assays to determine IC50 values for each of these three canonical substrates for SMR_Sa_ and SMR_Ft_. Mutation of the fully conserved, functionally critical glutamate residue in TM1 (E13Q-GdX, E14Q-EmrE, E13Q-SMR_Sa_, and E13Q-SMR_Ft_) renders the SMR transporter non-functional for either substrate binding or transport. Expression of this inactive variant serves as a negative control, ensuring that any difference in *E. coli* growth is due to EmrE activity and not the metabolic cost of membrane protein expression. Guanidinium resistance assays were performed in BW25113*ΔGdX* cells to ensure no interference from native GdX activity. Eth^+^ and MV^2+^ assays were performed in MG1655Δ*emrE* E. coli to ensure no interferences from native EmrE.

SMR_Sa_ and SMR_Ft_ are grouped into the EmrE-like SMR subfamily based on sequence similarity, and we therefore hypothesized that these SMRs would confer resistance to ethidium bromide and methyl viologen, but not guanidinium. To confirm this, we conducted SMR_Sa_ and SMR_Ft_ growth assays in the presence of guanidinium (Figure 2), ethidium, and methyl viologen (Figure 3). BW25113Δ*GdX E. coli* cells expressing either WT-GdX or transport-dead E13Q-GdX served as a positive control. Expression of functional GdX was required for cell growth, as expected, confirming that active GdX conferred resistance to guanidinium in this assay (Figure 2A). Repeating this experiment with SMR_Sa_ and SMR_Ft_ resulted in no growth when either WT or the transport dead E13Q variant was expressed, indicating that these SMR homologs do not confer resistance to guanidinium (Figure 2B, Supplemental Figure 3).

**Figure 2:**
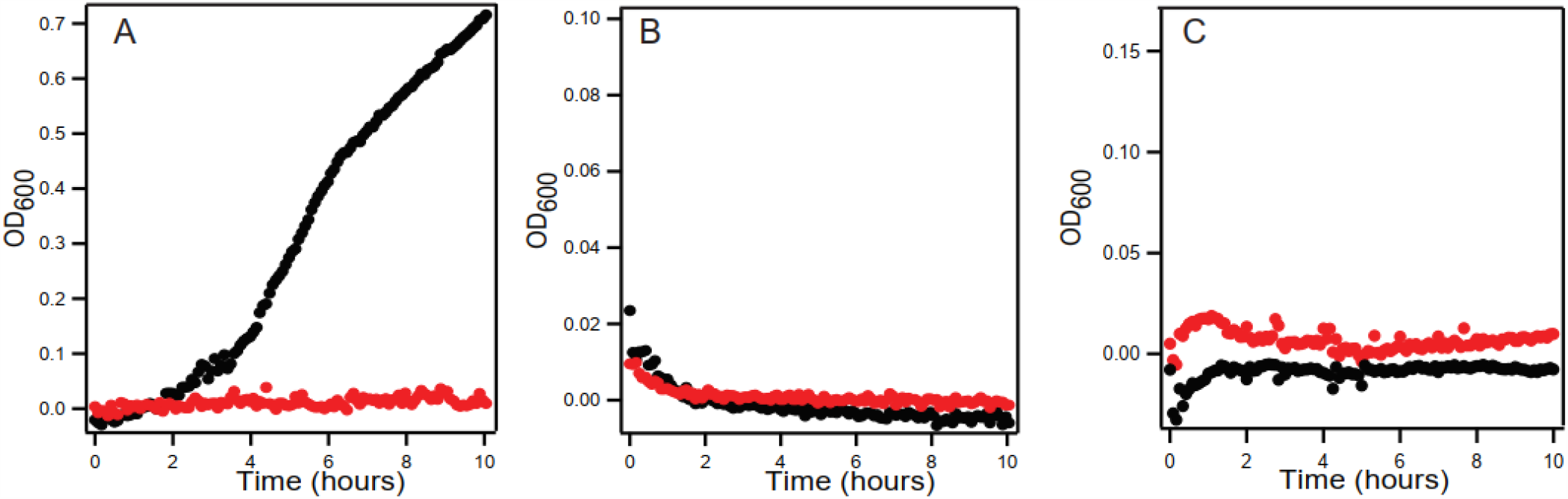
SMR_Sa_ and SMR_Ft_ do not confer resistance to guanidinium compared to GdX. Growth curves for MG1655 *ΔsugE E. coli* expressing GdX (A), SMR_Sa_ (B), and SMR_Ft_ (C) in the presence of 50mM guanidinium are displayed as OD_600_ versus time. Data for *E. coli* expressing functional, WT transporter is shown in black, and data for *E. coli* expressing non-functional E13Q mutants of each transporter is shown in red. Each point is the average of three technical replicates.

**Figure 3:**
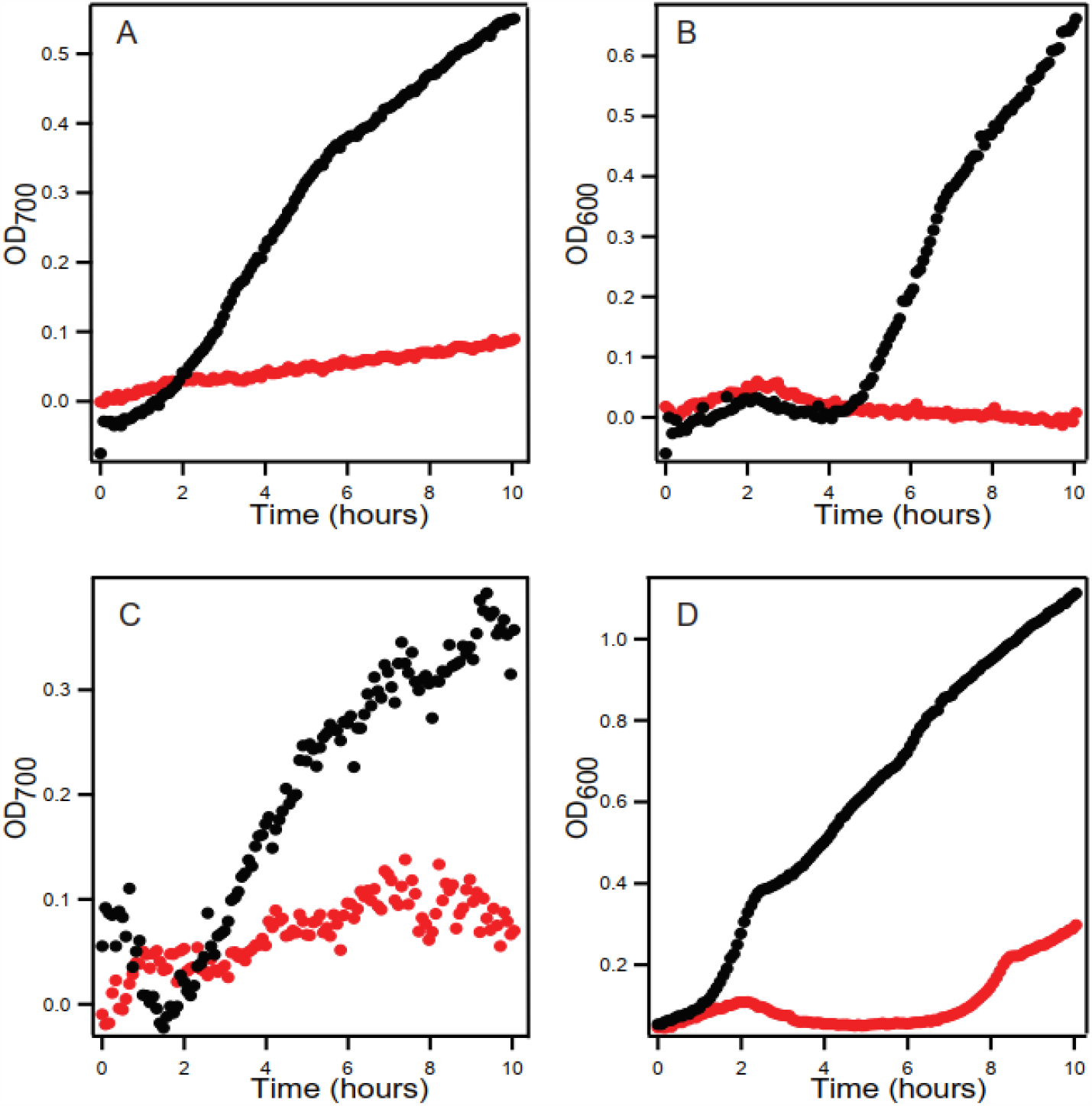
SMR_Sa_ and SMR_Ft_ confer resistance to ethidium bromide and methyl viologen. Growth curves for MG1655 *ΔemrE E. coli* expressing either SMR_Sa_ in the presence of 1mM ethidium bromide (A) or 0.3mM methyl viologen (B) or expressing SMR_Ft_ in the presence of 0.75mM ethidium bromide (C) of 0.3mM methyl viologen (D). Ethidium bromide results are shown as OD_700_ rather than OD_600_ to avoid the absorption spectrum of ethidium. Data for cells expressing WT SMR transporters are shown in black and data for cells expressing transport-dead mutants are shown in red. Each point is the average of three technical replicates.

To assess resistance to Eth^+^ and MV^2+^, we compared the growth of MG1655Δ*emrE E. coli* expressing either WT or the non-functional E13Q variant of each homolog. Eth^+^ and MV^2+^ are well-established EmrE substrates and are transported by many promiscuous multidrug pumps.

Representative growth curves of SMR_Sa_ and SMR_Ft_ are shown in Figure 3. Wild type SMR_Sa_ and SMR_Ft_ confer resistance to Eth^+^ (Figure 3, A and C; Supplemental Figure 1) that is comparable to resistance conferred by EmrE (11). The same is observed for MV^2+^, where MG1655Δ*emrE E. coli* expressing WT-SMR_Sa_ and cells expressing WT-SMR_Ft_ grow in the presence of 0.3mM MV^2+^, whereas the E13Q variants do not (Figure 3, B and D; Supplemental Figure 2). These results indicate that both SMR_Sa_ and SMR_Ft_ function similarly to EmrE *in vivo*.

### Biolog Functional Phenotyping MicroArrays reveal common compounds between SMRs

To broaden our understanding of the substrate specificity profiles for these transporters and to assess whether they only confer resistance or can confer both resistance and susceptibility in a substrate-specific manner, we performed Biolog Functional Phenotyping Microarrays. These microarrays are used to assess the bacterial metabolic output in the presence of a defined treatment: carbon source, nitrogen source, environmental salinity or pH, or a panel of drugs and small molecules (29). Here we assayed the chemical sensitivity panel, which includes many drug-like compounds of interest for understanding SMR activity. This panel of ten 96-well microplates includes 240 different compounds, with four wells are two different concentrations for each compound. Many known SMR substrates are polyaromatic cations, and this commercially available assay is designed to avoid interference from the natural fluorescence of these compounds, enabling a single consistent assay format for characterization across several hundred potential substrates.

We performed the Biolog microbial chemical sensitivity screen for EmrE, SMR_Sa_, and SMR_Ft._ Each screen is performed with two sets of plates side by side: one set with MG1655Δ*emrE E. coli* expressing WT functional transporter, and one set with MG1655Δ*emrE E. coli* expressing the corresponding non-functional transporter (E14Q-EmrE, E13Q-SMR_Sa_, and E13Q-SMR_Ft_). Thus, any difference observed in the metabolic output reflects the impact of SMR activity and not the metabolic impact of membrane protein expression. Metabolic activity is assayed through a colorimetric readout of NADH production over time for 24 hours, and the area under the curve represents the total metabolic activity. Higher metabolic activity by *E. coli expressing* WT transporter indicates that active transport is beneficial, consistent with the SMR transporter conferring resistance to the toxic compound in the well. In contrast, higher metabolic activity by *E. coli* expressing the transport-dead variant indicates that SMR activity is detrimental and the SMR transporter confers susceptibility rather than resistance.

The entire assay comparing functional and non-functional transporter is run in duplicate. The correlation between total metabolic output for the two replicates under each condition (WT or non-functional transporter), was consistently 0.85 (Supplemental Figures 4-6). For each screen, individual wells were compared between WT and non-functional transporter and scored as positive (resistance) or negative (susceptibility) hits if the difference above or below the 10% trimmed mean, as described in the methods. Each set of Biolog plates contains four wells at two different concentrations of a single compound. Since the entire assay was run twice for each transporter, this results in a total of 8 wells for a single compound, or a maximum possible score of ±8. To account for the non-zero rate of false-positives or false-negatives in scoring individual wells, as well as the potential that the transporter could confer resistance or susceptibility at one compound concentration but not the second concentration in the Biolog plates, we therefore selected any compound that scored ≥ +3 as a resistance hit and ≤ -3 as a susceptibility hit. All the known EmrE resistance substrates (methyl viologen and acriflavine) that are present in the Biolog compound set are selected as resistance hits using this scoring system, confirming the validity of this scoring method.

This analysis resulted in both susceptibility and resistance hits for all three SMR transporters in this study. When the data is sorted by the score of each compound for EmrE (Figure 4A), there is some correlation in compound scores and hits between homologs (clusters on the top right and bottom left of Figure 4A), but there are also hits unique to each transporter. The Venn diagram in Figure 4B illustrates the extent of these combined results among the proteins tested here. SMR_Sa_ had the highest number of unique results (12 total), with EmrE following (9 total) and SMR_Ft_ having the least (7 total). EmrE and SMR_Sa_ appear to have more compound hits in common (8 total) than EmrE shares with SMR_Ft_ (2 in total). Five compounds appeared as hits for all three homologs: methyl viologen, chelerythrine chloride, and cetylpyridinium chloride as resistance hits; and hexachlorophene and spectinomycin as susceptibility hits. Full results for the three transporters are included in the supporting Data Set.

**Figure 4:**
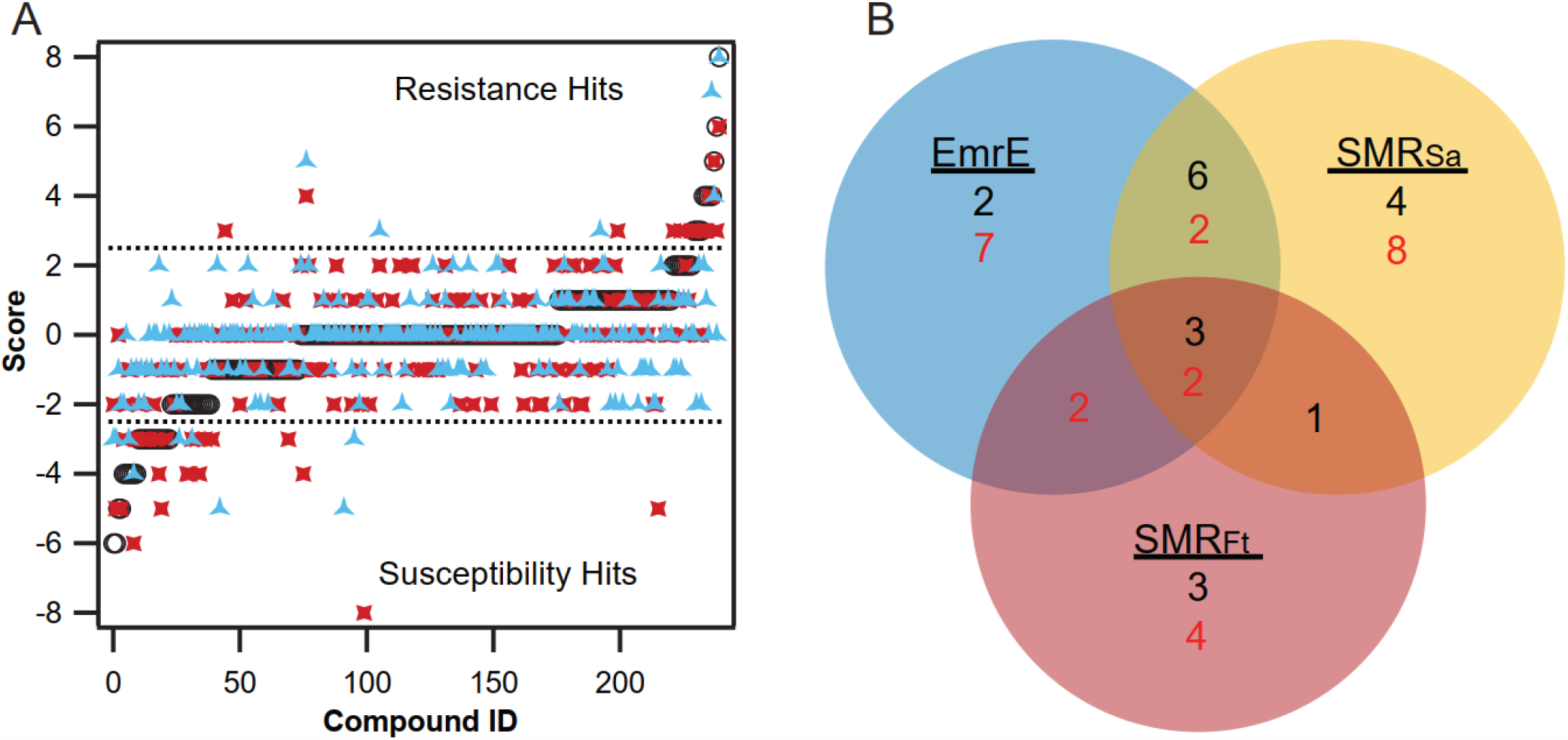
Comparison of Biolog microarray hits reveals similarities between SMR homolog hits. A) Overlaying the score of all three SMR homologs, sorted according to the EmrE scores for the compounds, reveals similarities in the strength of the resistance or susceptibility phenotype conferred by each homolog. EmrE results are depicted in black open circles, SMR_Sa_ in red squares, and SMR_Ft_ in blue triangles. The explanation of the scoring method can be found in the methods section. B) A Venn diagram showing the relationship between the hits for EmrE, SMR_Sa_, and SMR_Ft_. The number of resistance hits are listed in black and susceptibility hits are listed in red for individual transporters or are common hits across multiple transporters.

### Charge distinguishes resistance and susceptibility substrates of EmrE, SMR_Sa_, and SMR_Ft_

Previous studies of EmrE substrate specificity suggested that substrate charge and hydrophobicity were key parameters affecting the affinity and transport rate EmrE substrates (1, 11, 28). These studies focused on known compound classes to which EmrE confers resistance (polyaromatic cations, quarternary ammonium compounds, etc.). Using our larger Biolog dataset, we compared the hydrophobicity (cLogP) and predicted charge for all the hits for EmrE (Figure 5A), SMR_Sa_ (Figure 5B), and SMR_Ft_ (Figure 5C).

**Figure 5:**
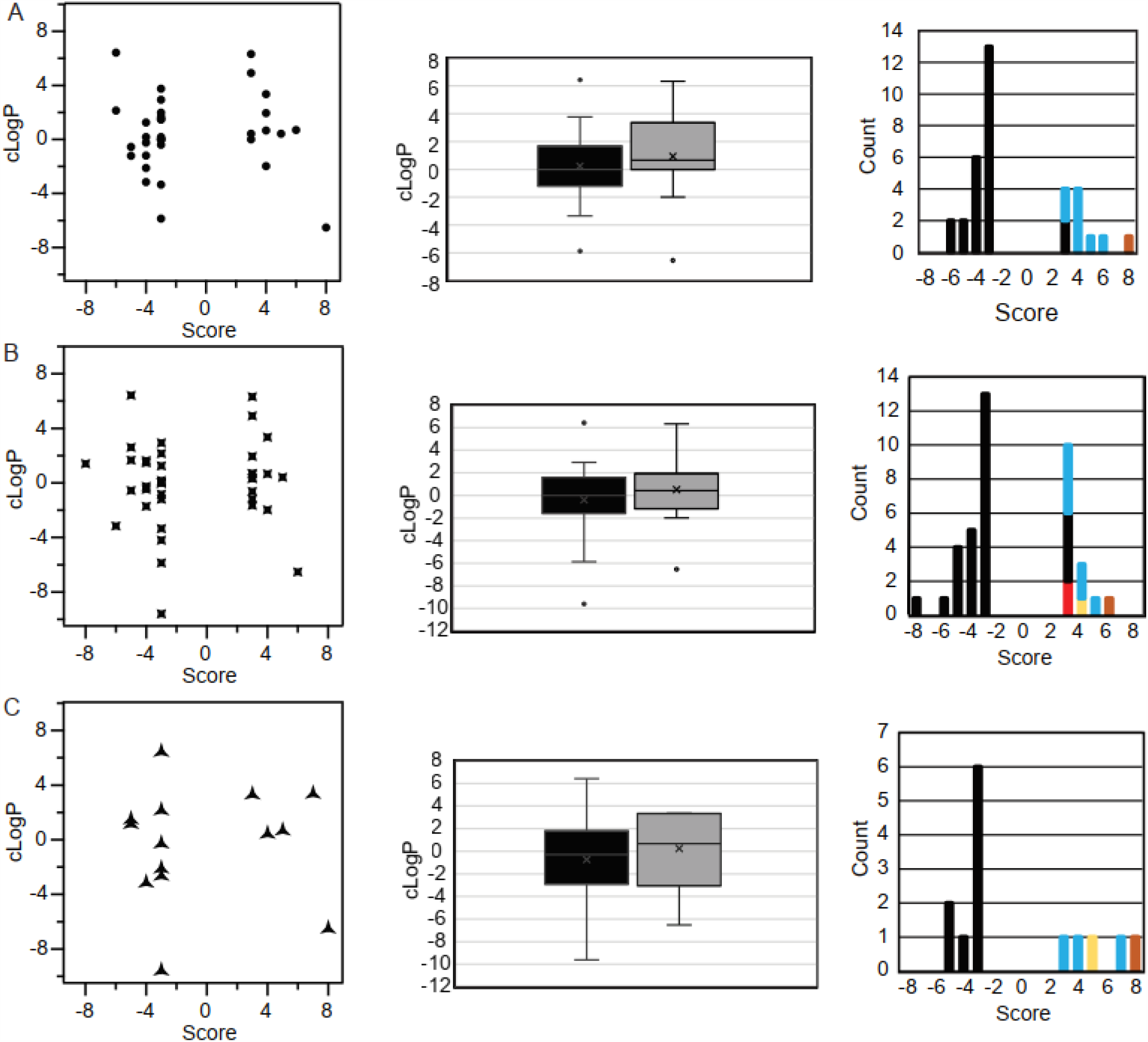
Hydrophobicity and charge of phenotypic microarray hits for EmrE, SMR_Sa_, and SMR_Ft_. Charts showing the cLogP and predicted formal charge of all hits for EmrE (A), SMR_Sa_ (B), and SMR_Ft_ (C). Susceptibility hits have a score of ≤ -3. Resistance hits have a score of ≤ 3. Box and Whisker plots of cLogP data were calculated using the standard quartile algorithm for each individual SMR. Susceptibility data is shown in black in the box and whisker plot. Resistance hit data is shown in grey in the box and whisker plot. Charge (third column) is plotted by score and count for each transporter. The color scheme is as follows: -2, red; -1, yellow; 0, black; 1, blue; and 2, rust.

Contrary to previous literature, hits from the Biolog microarray do not show any correlation between hydrophobicity and Biolog score for either susceptibility or resistance hits (first and middle column, Figure 5). However, charge does appear to distinguish resistance and susceptibility substrates of EmrE-like SMRs. Susceptibility hits for EmrE, SMR_Sa_, and SMR_Ft_ (third column, Figure 5) were universally uncharged at neutral pH. In contrast, the majority of resistance hits were positively charged, as expected based on previous literature (11). However, a few negatively charged or neutral resistance hits were also detected, and these will need to be confirmed, both as hits and the net charge of the substrates under the assay conditions. This trend is consistent across EmrE, SMR_Sa_, and SMR_Ft_ confirming the previous characterization that SMR transporters confer resistance to quaternary cation compounds, and that previous screens focusing on this compound class would have missed uncharged substrates to which SMR transporters confer susceptibility.

## DISCUSSION

Although SMR transporters are present across the bacterial kingdom and are known to confer broad toxin resistance, there is relatively little information about their substrate profiles. Beyond the founding members of each SMR subfamily, EmrE and GdX, there has been limited biochemical analysis of other SMR transporters, which have only been bioinformatically sorted into the EmrE-like family of SMRs or the GdX-like subfamily. Here, we experimentally confirm that SMRs from *S. aureus* (SMR_Sa_) and *F. tularensis* (SMR_Ft_), which have been classified as EmrE-like, match the expected resistance profile for this classification based on their ability to confer resistance to canonical substrates ethidium and methyl viologen (Figure 3) but not guanidinium (Figure 2B, C).

Previous work on EmrE shows that individual amino acid changes to the protein can alter the substrate profile (9, 25). Thus it is not unexpected that SMR homologs with different primary amino acid sequences would have a unique substrate profile that only partially overlaps with other SMR transporters. Biophysical data shows that EmrE also has mechanistic promiscuity, potentially performing uniport and symport (susceptibility) as well as the canonical antiport (resistance) activity (12, 27). Our Biolog data for EmrE show that it can confer resistance and susceptibility (Figure 4A) as predicted by the biophysical data. Furthermore, our results for SMR_Sa_ and SMR_Ft_ show that this mechanistic promiscuity extends beyond EmrE (Figure 4B). Although there is growing recognition that not all transporters are tightly coupled (30, 31), it is unusual to observe that the activity of a single transporter can result in opposing biological phenotypes for different substrates.

Prior substrate screens on EmrE have focused on compounds similar to traditional resistance substrates (quaternary ammonium cations, polyaromatic compounds, etc.) (1, 9, 11). With this less targeted panel of substrates from Biolog, we discovered greater substrate promiscuity and interesting differences in the characteristics of resistance and susceptibility substrates of EmrE-like SMRs.

Hydrophobicity was previously identified as an important substrate characteristic influencing substrate affinity and the rates of alternating access and transport by EmrE within a small set of tetraphenylphosphonium derivatives (9). However, our data reveals no correlation between hydrophobicity and resistance or susceptibility across the much broader set of substrates tested here (Figure 5, first and second column). In contrast, our results confirm the importance of charge in determining how a substrate is transported by EmrE-like SMRs. Previously, EmrE was characterized as a toxic cation antiporter, conferring resistance to quaternary ammonium compounds and polyaromatic cations. The resistance substrates identified in the Biolog assay are predominantly cationic and include the known EmrE substrates, consistent with this prior literature. However, the susceptibility substrates we have identified are uncharged. Previous work into EmrE substrates about charge only focused on cationic compounds and common classes of compounds transported by multidrug resistance transporters (11), which may be why these substrates have not been found before. Interestingly, metal ions were also identified as resistance hits for all three SMR transporters. This raises the question of whether EmrE or EmrE-like SMR transporters can transport metal ions directly, or perhaps transport metal ions bound by a siderophore, which would more closely resemble known SMR substrates. Thus, the broader Biolog screen reported here has expanded our knowledge of SMR transporter activity and promiscuity, and raises a number of interesting questions that require follow up with *in vitro* assays to better define the limits of both substrate and mechanistic promiscuity by SMR transporters.

## MATERIALS AND METHODS

### Bacterial Strains and Plasmids

MG1655 Δ*emrE E. coli* cells were used to assess EmrE-like functionality and BW25113 Δ*GdX* (32, 33) were used to assess GdX-like functionality *in vivo*. WT and variant (E13/14Q) versions of EmrE (AAN90039.1), GdX (AAC46453.1), *S. aureus* (WP_031824198) and *F. tularensis* (WP_003036323) SMRs were cloned into the pWB plasmid, a low copy, leaky-expression vector. These plasmids were used in all assays described in this paper.

### Microplate Growth Curves

Cells expressing plasmids of interest were grown in LB media (tryptone, yeast extract, 30mM bistris propane, 100µg/mL ampicillin, pH 7.0) from a single colony to an OD of 0.2. The cells were then diluted to a final OD of 0.01 in microplates (Corning, REF: 351172) containing concentration ranges of Eth^+^, MV^2+^, or guanidium. The plates were incubated in a microplate reader (BMG-Labtech) at 37°C. OD_600_ (guanidinium and methyl viologen) and OD_700_ (ethidium bromide) were measured every 5 minutes for 20 hours. Experiments were performed in technical and biological triplicate and data was analyzed using Excel and Igor Pro.

### Functional Phenotypic MicroArrays

MG1655 *ΔemrE E. coli* cells containing either WT- or E13Q-SMR_Xx_ constructs were grown on LB-Amp media overnight at 37°C. The phenotype microarray tests followed the established protocols of standard PM procedures for *E. coli* and other gram-negative bacteria (12). PM01-20 plates were used to screen both WT- and E13Q-SMR_Xx_ expressing cells (http://www.biolog.com/products-static/phenotype_microbial_cells_overview.php). Overnight plates were resuspended in IF-0a inoculating fluid (Biolog) to an optical density of 0.37. The cells were diluted by a factor of 6 into IF-0a media plus Redox Dye A and 20mM glucose was added for PM3-8 plates. Cells were diluted to a 1:200 dilution in IF-10a media (Biolog) with Redox Dye A for PM9-20 plates. PM plates were inoculated with 100µL of cell suspensions per well. The microplates were incubated at 37°C and read using the OmniLog instrument every 15min for 24 h. The area under the resulting metabolic curves was determined for cells expressing WT-SMR_Xx_ or E13Q-SMR_Xx_. The difference was calculated as

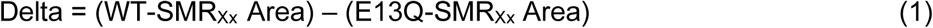

Resulting in positive values for greater respiration by cell expressing WT-SMR and negative values for greater respiration by cells expressing non-functional SMR. The 10% trimmed mean was then calculated for each data set (WT replicate 1, WT replicate 2, E13Q replicate 1, E13Q replicate 2) separately for each transporter as variation between replicates can arise due to minor deviations between plate sets or in the exact concentration of dye or OD of cells upon dilution on different days. The standard deviation was then calculated among known non-hits (selecting at least 50 wells out of the 960 total wells in a single data set) to determine the cut-off values for actual hits. Individual wells were assessed as hits if the calculated Delta value (equation 1) was more than two standard deviations from the 10% trimmed mean. For each hit, a value of +1 was assigned for resistance hits (positive Delta) and a value of -1 was assigned for susceptibility hits (negative Delta). These values were then summed across all eight wells for a single compound (4 wells of the same compound per plate set * 2 replicates, with a max score of ±8. Final resistance or susceptibility hits were assigned if the total score was ≥ +3 (resistance) or ≤ -3 (susceptibility). This definition was chosen since small total hit scores of ±1 or ±2 could arise by chance using the ± 2*SD cutoff to score individual wells. Values of ±3 recognize consistent hits across multiple replicates and/or different concentrations of the same compound. Our cutoff is not set higher since the 4 wells of each compound on a single plate set include different concentrations and some concentrations may not be sufficient to elicit a phenotype.

## ACKNOWLEDGEMENTS

The authors would like to thank Jason Peters for his intellectual contribution to this manuscript. The authors wish to thank Grant A. Hussey for aiding in the initial plasmid design of the pWB vector. Thanks also to Trey Sato for his assistance in using the OmniLog Instrument for Biolog Assays with funding from the Great Lakes Bioenergy Research Center under U.S. Department of Energy, Office of Science, Office of Biological and Environmental Research Award Number DE-SC0018409. This work was supported by the USDA National Institute of Food and Agriculture Hatch WIS01985. Research reported in this publication was supported by the institute for General Medical Sciences of the National Institutes of Health under award number R01GM095839. The content is solely the responsibility of the authors and does not necessarily represent the official views of the NIH, USDA, or NIFA. PJS was supported by a Departmental Fellowship from the Department of Biochemsitry at the University of Wisconsin-Madison.

## AUTHOR CONTRIBUTIONS

PJS performed Phenotypic MicroArrays and subsequent data analysis, wrote the manuscript, and supervised growth assays and IC50 assays. WFB performed growth assays and IC50 assays of EmrE and GdX, optimized these protocols for the other SMRs and edited the manuscript. BLY performed growth assays and IC50 assays of SMR_Sa_ and edited the manuscript. C JP performed growth assays and IC50 assays of SMR_Ft_ and edited the manuscript. KAHW supervised the project, aided in Biolog data analysis, edited the manuscript, and secured funding for the work.

